# Genome sequence of *Tacca chantrieri* reveals the genetic basis of floral pigmentation

**DOI:** 10.64898/2026.03.17.712415

**Authors:** Julie Anne V. S. de Oliveira, Boas Pucker

## Abstract

*Tacca chantrieri*, black bat flower, has showy flowers often appearing almost black. Here, we present the genome sequence and corresponding annotation to identify the genetic basis of the pigmentation. Candidate genes associated with the anthocyanin biosynthesis were identified based on this genome sequence and investigated with respect to their properties. The best dihydroflavonol 4-reductase (DFR) candidate, which harbours all amino acid residues believed to be required for DFR activity, shows a threonine in the substrate preference determining position where most characterized DFRs display asparagine or aspartate. This amino acid residue appears to be frequent in the Dioscoreaceae family as a comprehensive investigation revealed.

## Introduction

*Tacca chantrieri* (**Fig. 1**), also known as black bat flower, is a member of the yam family Dioscoreaceae with 10 species in the genus *Tacca* (Zhang *et al*., 2005). The species was first described by Édouard André in 1901 (Plants of the World, 2026). It inhabits regions with tropical, moist conditions. Despite the showy floral morphology, *T. chantrieri* is largely autonomous self pollinating (Zhang *et al*., 2006). This might be due to rare visits by pollinators or due to a previous interaction with a now extinct pollinator (Zhang *et al*., 2006). The dark color might serve as a carrion mimic in attracting flies (Zhang *et al*., 2005). *T. chantrieri* has been used in traditional Chinese medicine and there are reports suggesting health benefits (Tinley *et al*., 2003; Yang *et al*., 2020).

**Fig. 1.**
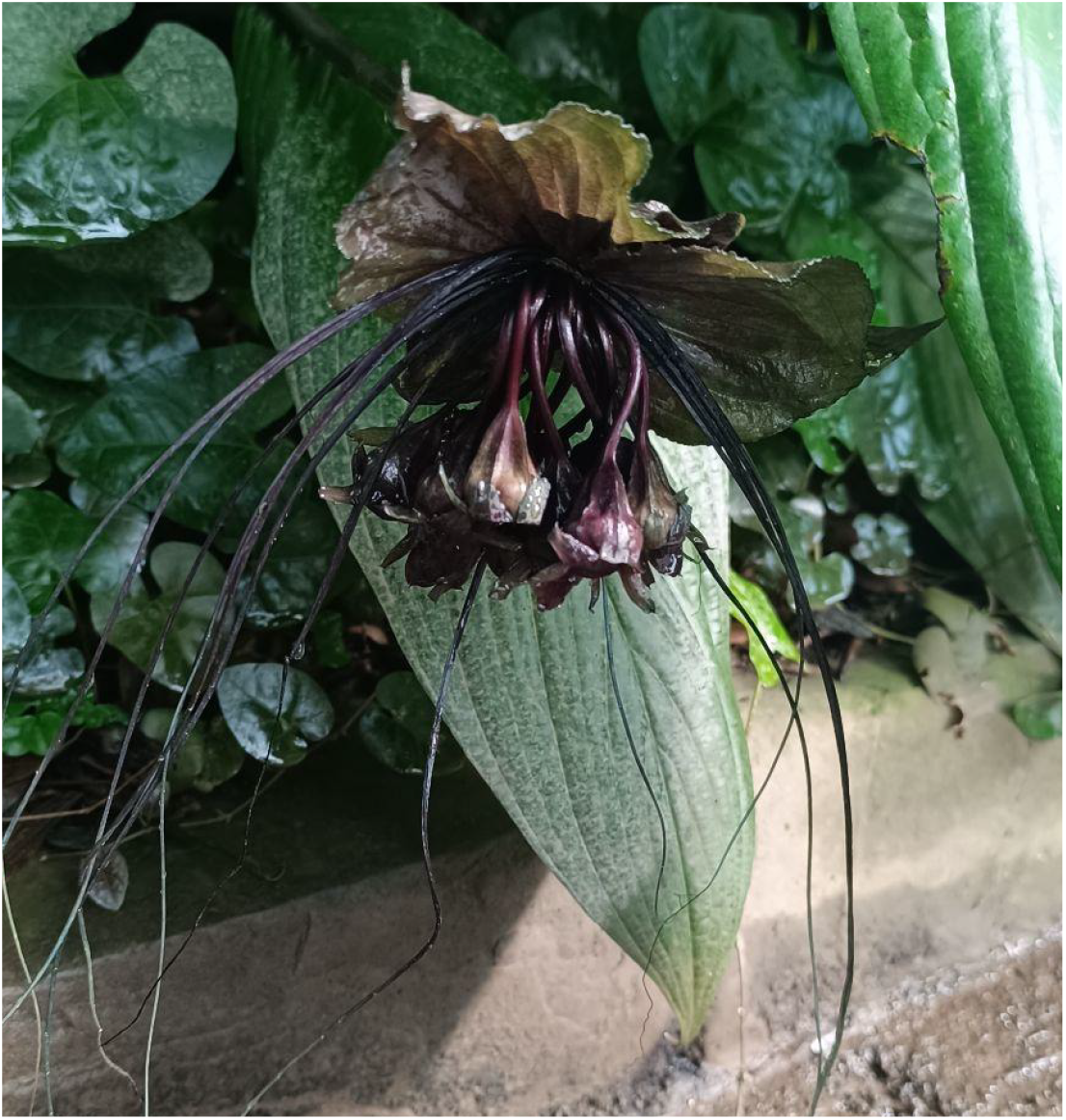
Picture of a dark pigmented flower of *Tacca chantrieri* (BONN-13679) taken in the tropical rainforest greenhouse of Bonn Botanical Gardens in July 2025.

The dark color of the showy *T. chantrieri* flower is likely due to accumulation of anthocyanins as these have been implicated in the coloration of many dark structures of various plant species (Wolff & Pucker, 2025). A range of different functions ranging from temperature benefits through passive heating to mimicry and camouflage have been proposed for dark pigmentation (Wolff & Pucker, 2025). Besides this dark pigmentation, anthocyanins can confer a number of different colors to plant structures including from orange, red, pink, and purple (Winkel-Shirley, 2001; Grünig *et al*., 2025). The ecological functions of anthocyanins include the attraction of pollinators and seed dispersers (Davies *et al*., 2012; Abid *et al*., 2022; Grünig *et al*., 2025) and the protection of leaves against strong light exposure (Gould *et al*., 2002; Gould, 2004; Merzlyak *et al*., 2008; Agati *et al*., 2021; Grünig *et al*., 2025). Due to their antioxidant properties, they are also believed to protect cell components under abiotic or biotic stress conditions (Gould, 2004; Kovinich *et al*., 2015; Naing & Kim, 2021; Jezek *et al*., 2023; Muralidhar *et al*., 2026).

Anthocyanins are produced through a branch of the flavonoid biosynthesis, which also leads to the formation of flavones, flavonols, and proanthocyanidins (Winkel-Shirley, 2001; Grotewold, 2006; Grünig *et al*., 2025). The precursor of these biosynthesis routes is phenylalanine that is channeled through the general phenylpropanoid pathway into the flavonoid biosynthesis (Winkel-Shirley, 2001). The biosynthesis of flavonoids is catalyzed by a number of well characterized enzymes in the core of the pathway and lineage-specific decorating enzymes (Grünig *et al*., 2025). Chalcone synthase (CHS), chalcone isomerase (CHI), and flavanone 3-hydroxylase (F3H) catalyze the first steps of the flavonoid biosynthesis that is shared by flavonol, anthocyanin, and proanthocyanidin branches (Forkmann *et al*., 1980; Ferrer *et al*., 1999; Jez *et al*., 2000). Diversity of the pathway is caused by potential hydroxylation of dihydrokaempferol, the F3H product, by flavonoid 3’-hydroxylase (F3’H) and flavonoid 3’,5’-hydroxylase (F3’5’H) (Seitz *et al*., 2007, 2015; Schwinn *et al*., 2014). Flavonol synthase (FLS) and dihydroflavonol 4-reductase (DFR) compete for the differently hydroxylated dihydroflavonols and channel metabolic flux towards flavonol or anthocyanin/proanthocyanidin biosynthesis, respectively (Johnson *et al*., 2001; Owens *et al*., 2008; Choudhary & Pucker, 2024). Amino acid residues at a specific position in DFR determine which of the three potential substrates is preferred (Johnson *et al*., 2001; Choudhary & Pucker, 2024). In many plant species, FLS and DFR show preferences for different dihydroflavonols and also an almost mutually exclusive expression pattern of the corresponding genes which could mitigate a direct competition (Choudhary & Pucker, 2024). Following DFR, anthocyanidin synthase (ANS), anthocyanin-related glutathione S-transferase (arGST), and UDP-dependent anthocyanidin 3-O-glucosyltransferase (U3GT) catalyze the next steps in the anthocyanin biosynthesis (Pelletier *et al*., 1997; Tohge *et al*., 2005; Eichenberger *et al*., 2023; Grünig *et al*., 2025). Several modifying enzymes can add additional sugar moieties, acyl groups, or methyl groups to anthocyanins leading to a huge variety of chemically different derivatives (Tohge *et al*., 2005; Luo *et al*., 2007; Yonekura-Sakakibara *et al*., 2012; Kovinich *et al*., 2014; Grünig *et al*., 2025).

Once synthesized at the endoplasmatic reticulum, anthocyanins are transported into the central vacuole, where they accumulate (Winkel-Shirley, 1999; Poustka *et al*., 2007; Kaur *et al*., 2021; Grünig *et al*., 2025). Two conceptually different transport routes have been proposed: (1) update into the ER and transfer via vesicles to the vacuole or (2) transport through the cytosol and direct import into the vacuole across the tonoplast (Goodman *et al*., 2004; Poustka *et al*., 2007; Kaur *et al*., 2021; Pucker & Selmar, 2022). It is possible that both routes exist and are active to different degrees in different plant lineages. While an initial model proposed the involvement of a GST protein as ligandin, i.e. protection of anthocyanins during transport through the cytosol, (Mueller *et al*., 2000; Conn *et al*., 2008), more recent evidence suggest that arGST might only be active as an enzyme (Eichenberger *et al*., 2023).

The complex biosynthesis of flavonoids including anthocyanins is controlled at the transcriptional level by numerous transcription factors (Weisshaar & Jenkins, 1998; Stracke *et al*., 2007; Gonzalez *et al*., 2008; Li, 2014; Xu *et al*., 2015; Grünig *et al*., 2025; Sielmann *et al*., 2026). Members of the MYB gene family play a particularly important role in the regulation of different branches of the flavonoid biosynthesis (Stracke *et al*., 2007, 2010; Li, 2014; Xu *et al*., 2015; Wheeler *et al*., 2022; Marin-Recinos & Pucker, 2024; Sielmann *et al*., 2026). Members of the MYB subgroup 7 (SG7) control the flavonol biosynthesis (Stracke *et al*., 2007, 2010). Members of the MYB subgroup 6 (SG6) and subgroup 5 (SG5) form a complex with bHLH and WD40 proteins to regulate the anthocyanin and proanthocyanidin biosynthesis, respectively (Nesi *et al*., 2000; Borevitz *et al*., 2000; Karppinen *et al*., 2021; Lafferty *et al*., 2022). This MBW transcription factor complex is a modular system which allows the replacement of one MYB component by another thus enabling specificity for different genes in the flavonoid biosynthesis (Ramsay & Glover, 2005; Li, 2014; Xu *et al*., 2015; Lloyd *et al*., 2017). It is assumed that the MYB and bHLH component bind to the DNA, while the WD40 protein (TTG1) acts as a scaffold protein holding MYB and bHLH together. Additional transcription factors have been implicated in the activation of anthocyanin biosynthesis in specific plant lineages (Lloyd *et al*., 2017; Grünig *et al*., 2025). Besides a large number of activators, there are also negative regulators that can interfere with the MBW complex or block cis-regulatory elements in the promoters of target genes (LaFountain & Yuan, 2021).

Here, we report about a genome sequence and corresponding annotation of *T. chantrieri* which revealed the genetic basis underlying the dark flower coloration. Candidate genes encoding enzymes and transcription factors associated with the anthocyanin biosynthesis are presented. An in-depth inspection of the DFR candidate polypeptide sequences revealed threonine at the substrate preference determining position. Given that this residue has been observed in multiple plant lineages above the species level, this might represent an adaptive change rather than a spontaneous variant.

## Materials and Methods

### DNA extraction and nanopore sequencing

DNA samples were taken from BONN-13679, which is cultivated in a tropical greenhouse in the Botanical Gardens of the University of Bonn. Temperature is maintained at 24°C during the day, and at 20°C during the night, with a humidity varying from 70% to 80%. Samples for DNA extraction were taken in July 2025. DNA extraction from young leaves was conducted following an established CTAB-based protocol for high molecular weight DNA extraction from plants (Siadjeu *et al*., 2020). The general sequencing and assembly workflow followed the guidelines provided previously (de Oliveira *et al*., 2026). In brief, high molecular DNA was subjected to library preparation with the SQK-LSK114 kit (Oxford Nanopore Technologies). Sequencing was conducted on R10 flow cells on a PromethION 2 Solo. Basecalling was conducted on a GPU in the de.NBI cloud using dorado v1.1.1 (Oxford Nanopore Technologies).

### Genome sequence assembly

Different assemblers were applied to obtain the best possible genome sequence. Shasta v0.14.0 (Shafin *et al*., 2020) was run on HERRO-corrected reads with the model Nanopore-r10.4.1_e8.2-400bps_sup-Herro-Jan2025 and using a minimal read length cutoff at 10kb. NextDenovo v2.5.2 (Hu *et al*., 2024) was also run with a read length cutoff at 10kb, an estimated genome size of 420 Mbp, and additional settings provided via a config file (Additional file B).

Hifiasm v-0.25.0-r726 (Cheng *et al*., 2021) was run on uncorrected ONT reads with default parameters as it performs its own internal read correction. The Hifiasm output was processed with the gfa2fa function of gfatools to obtain the final assembly in FASTA format. Basic statistics of all assemblies were calculated with contig_stats.py following a previously established workflow to assess the contiguity (de Oliveira *et al*., 2026). Assembly completeness was evaluated with BUSCO v6.0.0 (Simão *et al*., 2015; Tegenfeldt *et al*., 2025) using default parameters and the liliopsida_odb12 lineage.

### Prediction of protein encoding genes

GeMoMa v1.9 (Keilwagen *et al*., 2019) was run with the parameters pc=true pgr=true p=true o=true GAF.f=“start==‘M’ and stop==‘*’ and (score/aa>=‘0.75’)”. Hints were obtained from *Dioscorea zingiberensis* (GCA_026586065.1), *Dioscorea sansibarensis* (GCA_052858535.1), *Dioscorea alata* (GCA_020875875.1), *Wolffia australiana* (GCF_029677425.1), *Zostera marina* (GCA_001185155.1), *Spirodela intermedia* (GCA_902729315.2, GCA_902703425.1), and *Colocasia esculenta* (GCA_009445465.1). In addition, RNA-seq reads (SRR19179844) were obtained from the Sequence Read Archive with fastq-dump (NCBI, 2020; Katz *et al*., 2022) and mapped to the genome sequence with HISAT2 v2.2.1 (Kim *et al*., 2019) using default parameters to provide additional hints in the gene prediction. First filtering of the GeMoMa output was conducted with f=“start==‘M’ and stop==‘*’ and (score/aa>=‘0.75’)”. Next, RNA-seq evidence was prepared with GeMoMa’s ERE based on the read mapping with HISAT2. Attributes of external annotations were added with AnnotationEvidence. Filtering and merging of annotations was done with GeMoMa GAF: f=“start==‘M’ and stop==‘*’ and aa>=30 and avgCov>0 and (isNaN(bestScore) or bestScore/aa>=2.5) and iAA>=0.5 and pAA>=0.5” atf=“iAA>0.9 and pAA>0.9 and sumWeight>1 and avgCov>100 and tpc==1 and tie==1”. Finally, predicted genes were renamed with the AnnotationFinalizer and CDS and polypeptide sequences were extracted with GeMoMa’s Extractor.

BRAKER v3 (Gabriel *et al*., 2024) was run on the genome sequence with hints from the same RNA-seq read mapping. In addition, polypeptide sequences of viridiplantae and sequences obtained from UniProt were provided as external hints (Tegenfeldt *et al*., 2025; The UniProt Consortium, 2025).

The completeness of predicted sets of protein encoding genes was assessed with BUSCO v6.0.0 (Tegenfeldt *et al*., 2025) based on the liliopsida_odb12 data set and default parameters.

### Functional annotation of protein encoding genes

A general prediction of gene functions was conducted with construct_anno.py based on Araport11 information (Cheng *et al*., 2017; de Oliveira *et al*., 2026). Structural genes of the flavonoid biosynthesis were identified via KIPEs v3.2.7 (Rempel *et al*., 2023) using the flavonoid biosynthesis bait and residue set v3.4. Transcription factors of the MYB and bHLH gene family were annotated with the MYB_annotator v1.0.3 (Pucker, 2022) and bHLH_annotator v1.04 (Thoben & Pucker, 2023), respectively. TTG1 was identified based on orthology to previously characterized TTG1 genes in other plant species. Candidate sequences were identified with collect_best_BLAST_hits.py (Pucker & Iorizzo, 2023) and aligned with the bait sequences using MAFFT v7.505 (Katoh & Standley, 2013) and default parameters. The alignment was trimmed with algntrim.py (Pucker & Iorizzo, 2023) and subjected to IQ-TREE v1.6.12 (Nguyen *et al*., 2015) for inference of a tree. iTOL v6 (Letunic & Bork, 2024) was utilized for the visualization of the resulting tree.

For in-depth analyses of the MYB and bHLH candidates, alignments were generated with MAFFT v7.505 (Katoh & Standley, 2013) and gene trees were inferred with IQ-TREE v1.6.12 (Nguyen *et al*., 2015). Information for the inspection of residues in the DFR candidates was retrieved from previous studies about the substrate specificity of DFR types (Johnson *et al*., 2001; Miosic *et al*., 2014; Choudhary & Pucker, 2024). Information about the crucial residues for the interaction of MYB and bHLH proteins in the MBW complex was obtained from previously generated compilation (Pucker *et al*., 2024).

### Analysis of Dioscoreaceae DFR

The predicted polypeptide sequences of different Dioscoreaceae species including GCA_026586065.1 (Li *et al*., 2022), GCA_020875875.1 (Bredeson *et al*., 2022), GCA_052858535.1 (*Dioscorea sansibarensis*), and GCF_009730915.1 (*Dioscorea cayenensis* subsp. *rotundata*) were retrieved from the NCBI. Additional genome sequences of GCA_051820375.1 (*Dioscorea alata*), GCA_002260645.1 (*Dioscorea rotundata*), GCA_002260665.1 (*Dioscorea cayenensis* subsp. *rotundata*), GCA_014060945.1 (*Dioscorea zingiberensis*), GCA_051820395.1 (*Dioscorea alata*), GCA_902712375.1 (*Dioscorea dumetorum*, (Siadjeu *et al*., 2020)), GCA_005019695.1 (*Trichopus zeylanicus* subsp. *travancoricus*), and GCA_965111675.1 (*Dioscorea polystachya*), were retrieved from the NCBI annotated with GeMoMa following the workflow described above using the annotation of GCF_009730915 as hints. KIPEs v3.2.7 (Rempel *et al*., 2023) was run on all sets of polypeptide sequences to identify DFR candidates. A collection of DFR candidates from all species with >90% of functionally important amino acid residues was compiled and subjected to MAFFT v7.505 (Katoh & Standley, 2013) for construction of an alignment. A manual alignment inspection was conducted with seaview (Gouy *et al*., 2021). The alignment was trimmed with algntrim.py and default parameters. IQ-TREE v1.6.12 (Nguyen *et al*., 2015) was applied to infer a gene tree.

## Results & Discussion

### *Tacca chantrieri* genome sequence and structural annotation

Different assemblies were generated with Shasta, NextDenovo, and Hifiasm to obtain the best possible genome sequence of *Tacca chantrieri* (Table 1). Based on the assembly size of 540 Mbp and high contiguity indicated by the N50 of 31.4 Mbp, the Hifiasm assembly was selected as the representative genome sequence. All downstream analyses were conducted based on this genome sequence and all related datasets are available via bonndata (Vieira Salgado de Oliveira & Pucker, 2026). The size of the *T. chantrieri* genome sequence is in the range of previously reported genome sequences of *Dioscoreaceae* species (Tamiru *et al*., 2017; Siadjeu *et al*., 2020; Waweru *et al*., 2024).

**Table 1.**
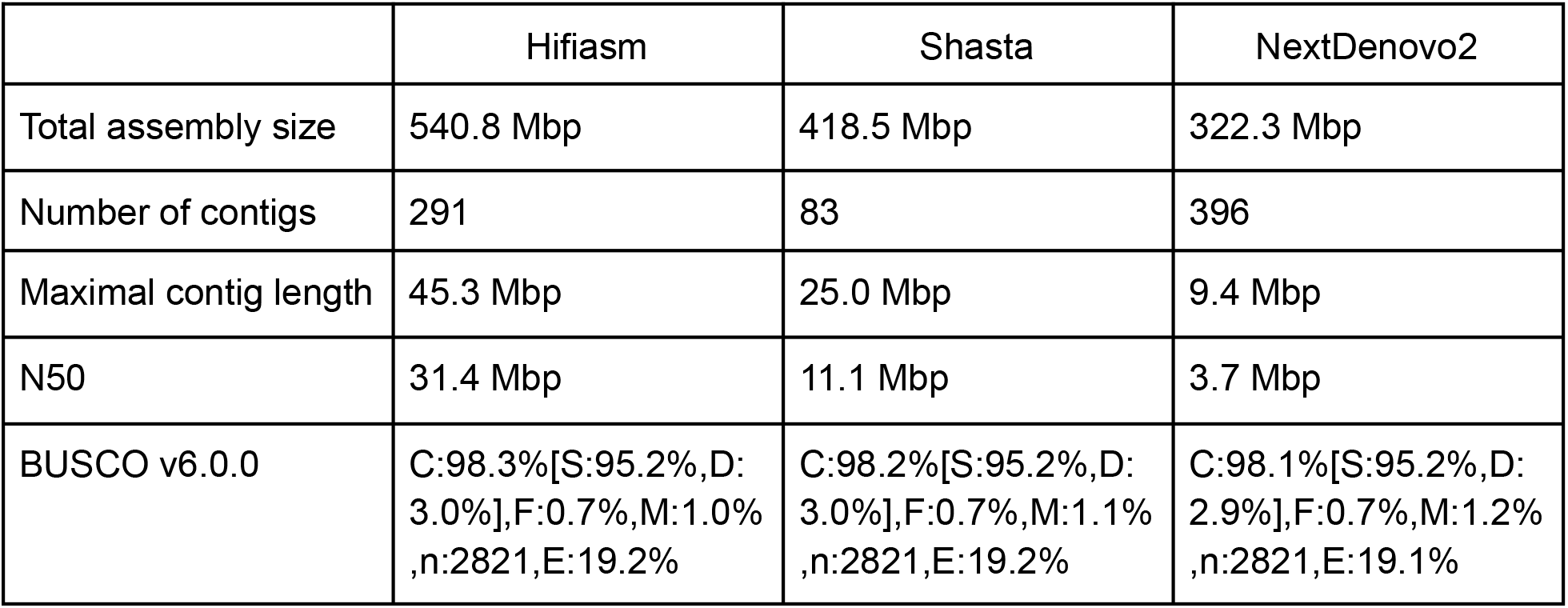
Statistics of the *Tacca chantrieri* genome sequence produced in this study. Assemblies have been trimmed by removing contigs <100 kb before statistics calculation. BUSCO was run with the liliopsida_odb12 dataset.

|  | Hifiasm | Shasta | NextDenovo2 |
| --- | --- | --- | --- |
| Total assembly size | 540.8 Mbp | 418.5 Mbp | 322.3 Mbp |
| Number of contigs | 291 | 83 | 396 |
| Maximal contig length | 45.3 Mbp | 25.0 Mbp | 9.4 Mbp |
| N50 | 31.4 Mbp | 11.1 Mbp | 3.7 Mbp |
| BUSCO v6.0.0 | C:98.3%[S:95.2%,D:3.0%],F:0.7%,M:1.0%,n:2821,E:19.2% | C:98.2%[S:95.2%,D:3.0%],F:0.7%,M:1.1%,n:2821,E:19.2% | C:98.1%[S:95.2%,D:2.9%],F:0.7%,M:1.2%,n:2821,E:19.1% |

Protein encoding genes were predicted in the representative genome sequence of *T. chantrieri* via BRAKER3 and GeMoMa to obtain the best possible annotation (Table 2). Both annotation approaches utilized the available RNA-seq data sets as hints. BRAKER3 was supported with protein sequences from other plant species as hints, while GeMoMa was supplied with the annotation of several related plant species to derive hints. Based on the higher number of complete BUSCO genes, the GeMoMa annotation was selected as the representative structural annotation. All downstream analyses exploring gene functions are based on this annotation. The GFF file and extracted coding and polypeptide sequences are available via bonndata (Vieira Salgado de Oliveira & Pucker, 2026). The number of predicted protein encoding genes is in the range previously reported for other plant species (Pucker & Brockington, 2018) and also aligns with the gene number reported for the closely related *Dioscorea dumetorum* genome sequence (Siadjeu *et al*., 2020).

**Table 2.**
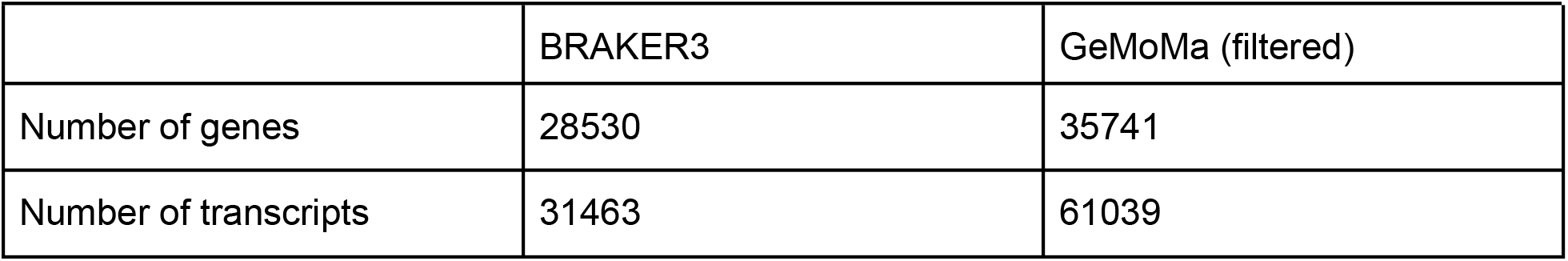
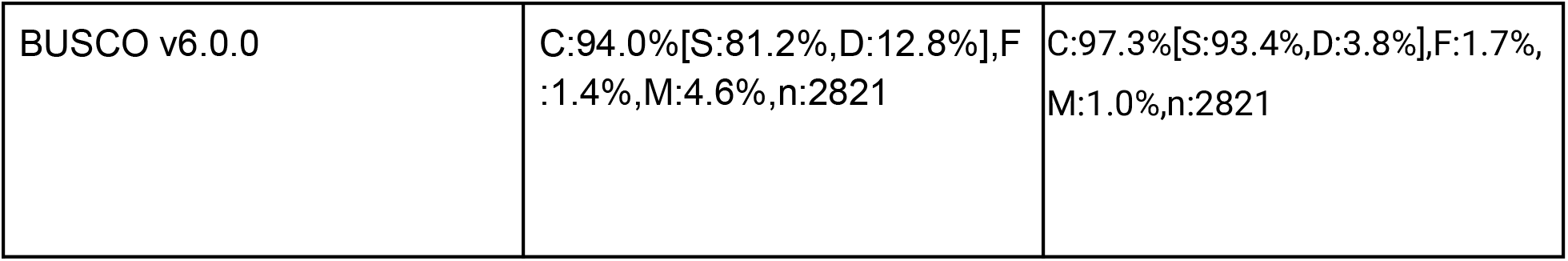
Comparison of different structural annotation approaches of *Tacca chantrieri*. BRAKER3 was run with RNA-seq hints and protein hints of viridiplantae. GeMoMa was supplied with RNA-seq hints and hints generated from several plant species (see methods).

|  | BRAKER3 | GeMoMa (filtered) |
| --- | --- | --- |
| Number of genes | 28530 | 35741 |
| Number of transcripts | 31463 | 61039 |
| BUSCO v6.0.0 | C:94.0%[S:81.2%,D:12.8%],F:1.4%,M:4.6%,n:2821 | C:97.3%[S:93.4%,D:3.8%],F:1.7%,M:1.0%,n:2821 |

### Structural genes in the anthocyanin biosynthesis of *Tacca chantrieri*

Genes of the flavonoid biosynthesis, including anthocyanin biosynthesis genes, were identified via KIPEs based on the structural annotation (**Fig. 2**, Additional file A). Several best candidates including CHI, ANS, F3’H, and F3’5’H show a deviation from the pattern of highly conserved amino acid residues in positions considered relevant for the enzyme function. Given that *T. chantrieri* is apparently able to produce high levels of anthocyanins, this suggests that lineage-specific differences exist in the Dioscoreaceae. Numerous studies on closely related yam species (Dioscorea) indicated that cyanidin derivatives account for most of the anthocyanin content (Wang *et al*., 2024; Chen *et al*., 2025; Zhang *et al*., 2025; Sun *et al*., 2025).

**Fig. 2.**
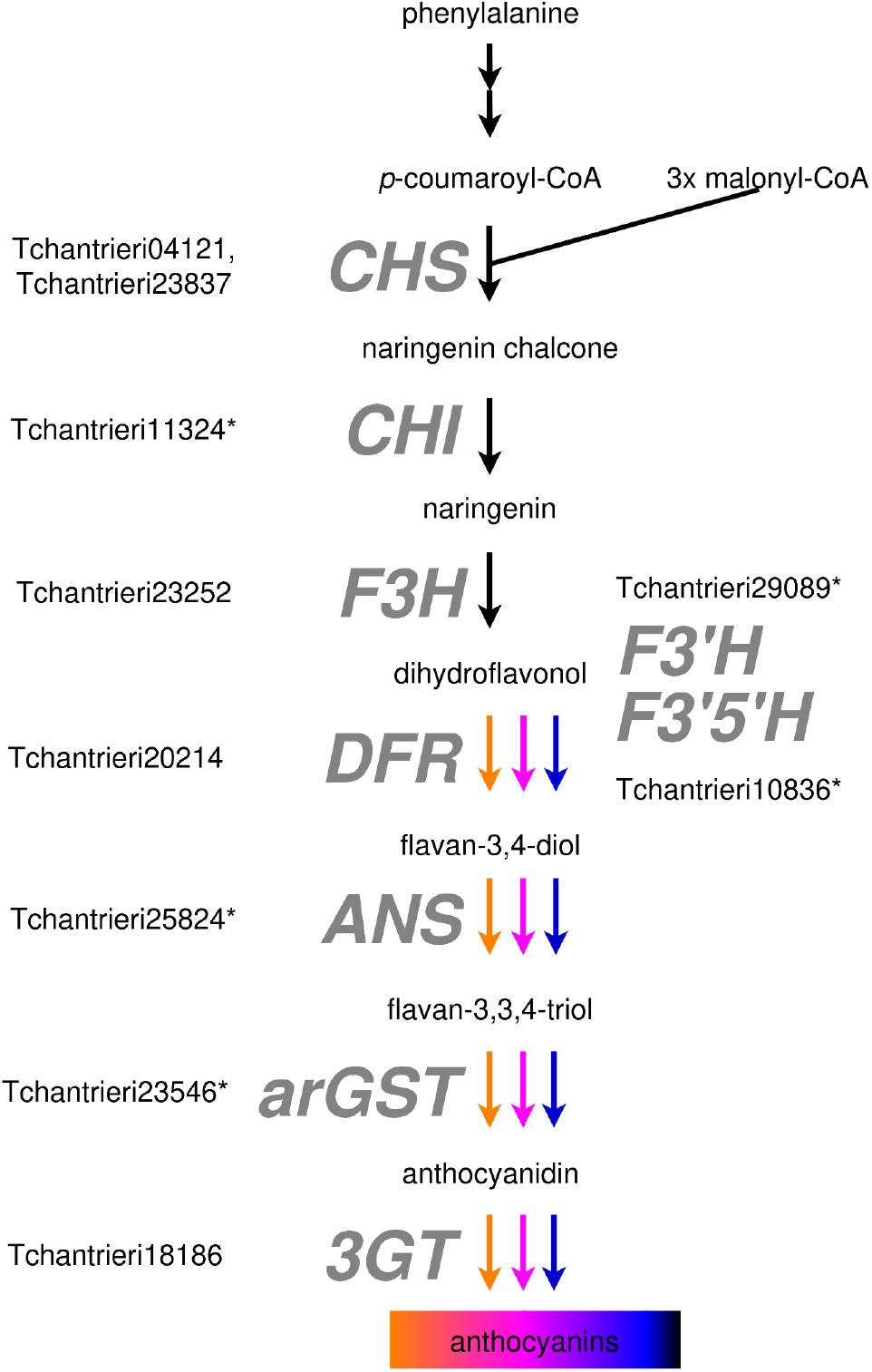
Structural genes of the anthocyanin biosynthesis. Candidates lacking an important amino acid residue are marked with an asterisk. CHS, chalcone synthase; CHI, chalcone isomerase; F3H, flavanone 3-hydroxylase; F3’H, flavonoid 3’-hydroxylase; F3’5’H, flavonoids 3’,5’-hydroxylase; DFR, dihydroflavonol 4-reductase; ANS, anthocyanidin synthase; arGST, anthocyanin-related glutathione S-transferase; and 3GT, UDP-dependent anthocyanidin 3-O-glucosyltransferase.

The best DFR candidates in *T. chantrieri* were inspected with respect to a diagnostic amino acid residue that is crucial for determining substrate preferences (Johnson *et al*., 2001; Choudhary & Pucker, 2024). The best DFR candidate showing all amino acid residues considered crucial for the enzymatic activity displays a threonine in the position where usually an asparagine or aspartate should be located (**Fig. 3**). A well characterized exception from the situation in most other plants is strawberry, which harbors a DFR copy that displays alanine in the substrate preference determining position (Miosic *et al*., 2014). As a consequence, this strawberry DFR copy has a high preference for dihydrokaempferol instead of dihydroquercetin (Miosic *et al*., 2014). Other deviations from the canonical N/D at this position were cystein in *Musa acuminata*, alanine in *Spirodela polyrrhiza*, and threonine in *Dioscorea alata* and *D. rotundata* (Choudhary & Pucker, 2024). A phylogenetic investigation of the DFR candidates in *T. chantrieri* supported the previously observed split into two lineages (Additional file C). To further corroborate the identity of the best DFR candidate and to explore the extent of T at the substrate preference determining position, an analysis across Dioscoreaceae was conducted. There are two DFR lineages in Dioscoraceae that display T (red, blue), while another DFR lineage shows N (yellow) (**Fig. 3c**). The emergence of non-canonical DFR versions was previously associated with ‘false organs’ i.e. a particular organ function taken over by a structure with an originally different organ identity like the ‘false fruit’ in strawberry or ‘false petal’ in *Anthurium andraeanum* (Choudhary & Pucker, 2024).

**Fig. 3.**
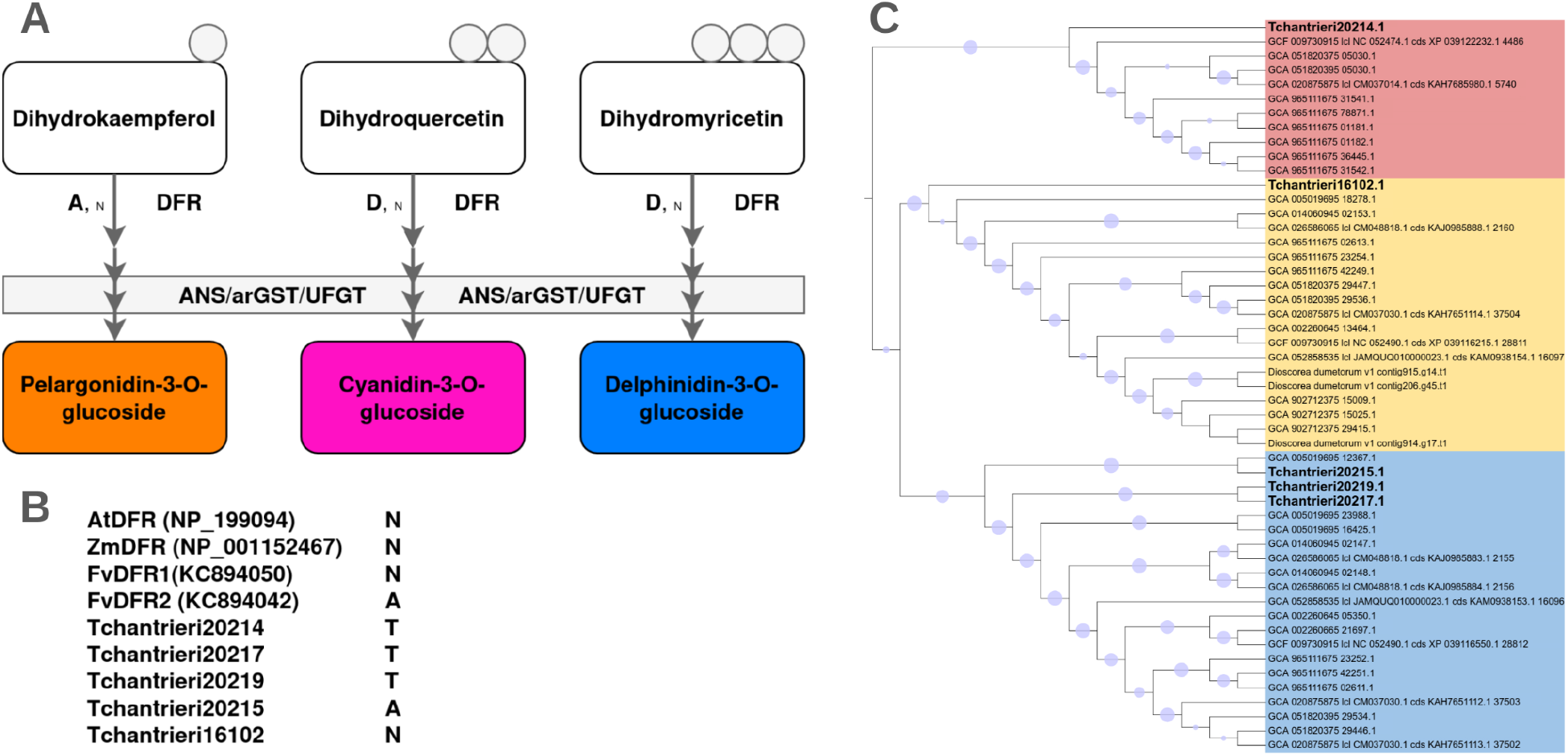
Investigation of the DFR type in *T. chantrieri*. Substrate preferences of different DFR types depend on amino acid residue at highlighted position (A) based on Johnson et al., 2001 and Choudhary & Pucker, 2024. Inspection of amino acid residues at substrate preference determining position (B). Phylogenetic analysis of DFR in the Dioscoraceae showing three distinct DFR clades (C).

### Transcriptional regulation of the anthocyanin biosynthesis in *Tacca chantrieri*

Transcription factors associated with the anthocyanin biosynthesis were identified based on orthology to previously characterized transcription factors in other plant species (**Fig. 4**, Additional file A). There are four copies of the anthocyanin-specific MYB activators (Tchantrieri00154, Tchantrieri04063, Tchantrieri00152, Tchantrieri15529), one copy of the corresponding bHLH partner (Tchantrieri07382), and two TTG1 candidates (Tchantrieri25454, Tchantrieri04146). The four MYB candidates are close paralogs of each other suggesting that there is only one active PAP lineage in *Tacca*. The importance of transcription factors for the activation of the anthocyanin biosynthesis has recently been investigated in the closely related genus *Dioscorea* by comparison of anthocyanin-pigmented and unpigmented yam species (Wang *et al*., 2024; Chen *et al*., 2025; Zhang *et al*., 2025). The large numbers of differentially expressed genes reported in these studies suggest that the causal factor for the difference is a variation in a transcription factor as observed before in numerous other plant species (Marin-Recinos & Pucker, 2024).

**Fig. 4.**
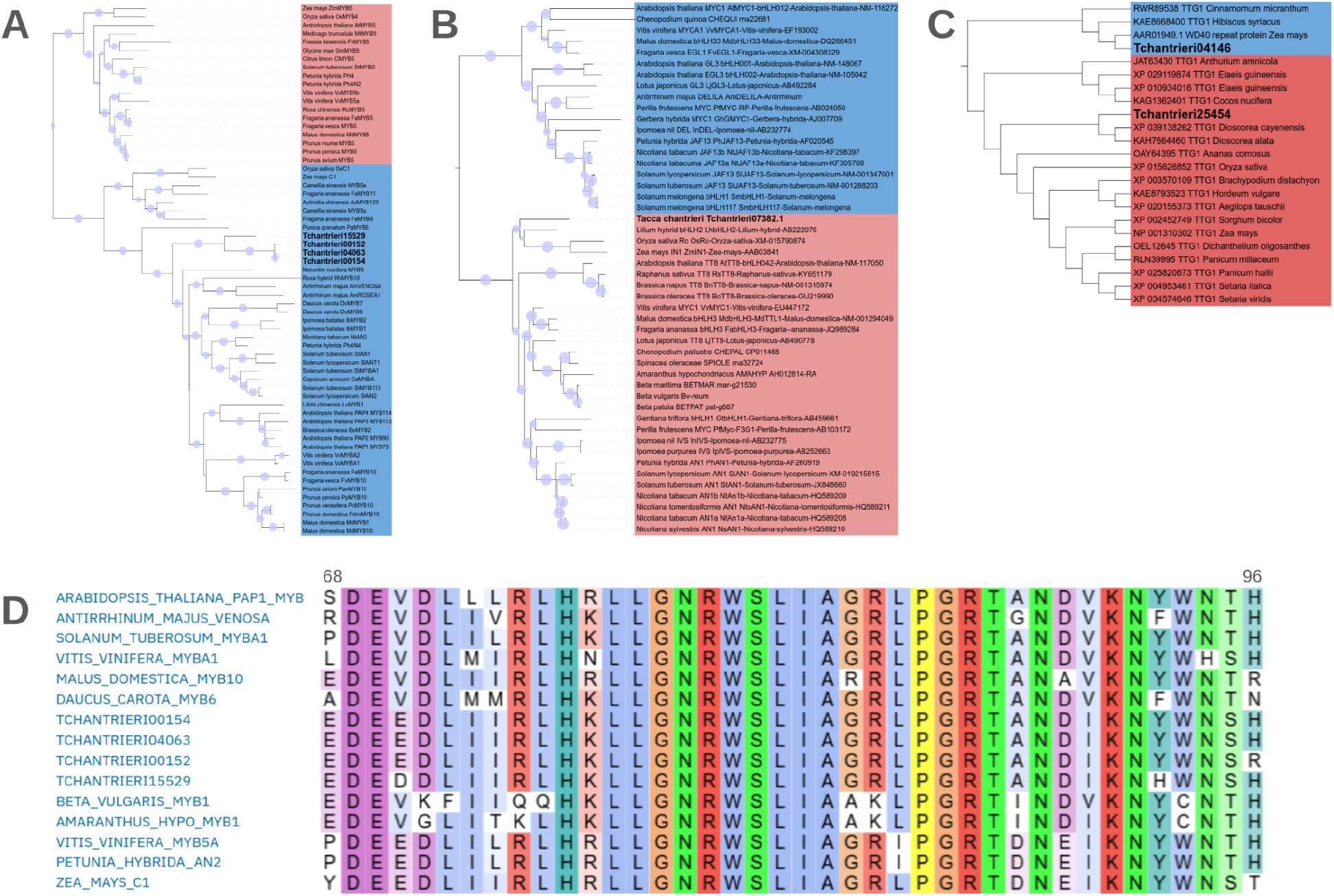
Investigation of the transcription factors involved in the regulation of the anthocyanin biosynthesis. (A) MYB subgroup 6 members, (B) bHLH transcription factors of the JAF13/TT8 lineage, (C) TTG1 candidates, and (D) inspection of the MYB-bHLH interaction domain check. Circles on the trees symbolize the bootstrap values. Positions in the alignment of MYB sequences are based on the *A. thaliana* MYB75.

The interaction of MYB and bHLH proteins for the formation of the MBW complex depends on a bHLH-interaction domain in the MYB protein that displays highly conserved residues (Zimmermann *et al*., 2004; Hatlestad *et al*., 2015; Sakuta *et al*., 2021; Pucker *et al*., 2024). Since most of these interactions have been investigated in the eudicots, we screened the *T. chantrieri* candidate sequences for monocot specific deviations (**Fig. 4D**). A comparison against MYB sequences of the betalain-pigmented sugar beet and amaranth served as negative control, as these MYBs have been co-opted for betalain biosynthesis regulation and are believed to have lost their ability to form MBW complexes (Hatlestad *et al*., 2015; Pucker *et al*., 2024).

## Conclusions

*Tacca chantrieri* is famous for its black flowers which inspired some of the trivial names. With the genome sequence and corresponding annotation presented here, it was possible to identify the structural gene and transcription factors candidates in the anthocyanin biosynthesis, which influence floral color. This provides a molecular basis for future studies addressing the ecological relevance of flower morphology and coloration. The substrate preferences of DFR with a threonine in the determining position offer potential for future discoveries. This could be particularly relevant in the context of pollinator attraction or the lack of such plant-insect interactions.

## Declarations

### Ethics approval and consent to participate

Not applicable

### Consent for publication

Not applicable

### Availability of data and materials

All datasets associated with the *Tacca chantrieri* genome sequence and annotation are available via bonndata (https://doi.org/10.60507/FK2/6SOMER).

### Authors’ contributions

JAVSdO conducted the DNA extraction, sequencing, and bioinformatic analyses. BP supervised the work, conducted bioinformatic analyses, and wrote the manuscript. All authors approved the final version of the manuscript and agreed to its submission.

## Acknowledgements

This work was supported by the de.NBI Cloud within the German Network for Bioinformatics Infrastructure (de.NBI) and ELIXIR-DE (Forschungszentrum Jülich and W-de.NBI-001, W-de.NBI-004, W-de.NBI-008, W-de.NBI-010, W-de.NBI-013, W-de.NBI-014, W-de.NBI-016, W-de.NBI-022). We thank all members of the Plant Biotechnology and Bioinformatics group for their support and feedback during the process. We are grateful for the excellent support provided by the team of the University of Bonn Botanic Gardens.

