## Supplementary figures and images for "Genome sequence of *Tacca chantrieri* reveals the genetic basis of floral pigmentation"

### Additional file C

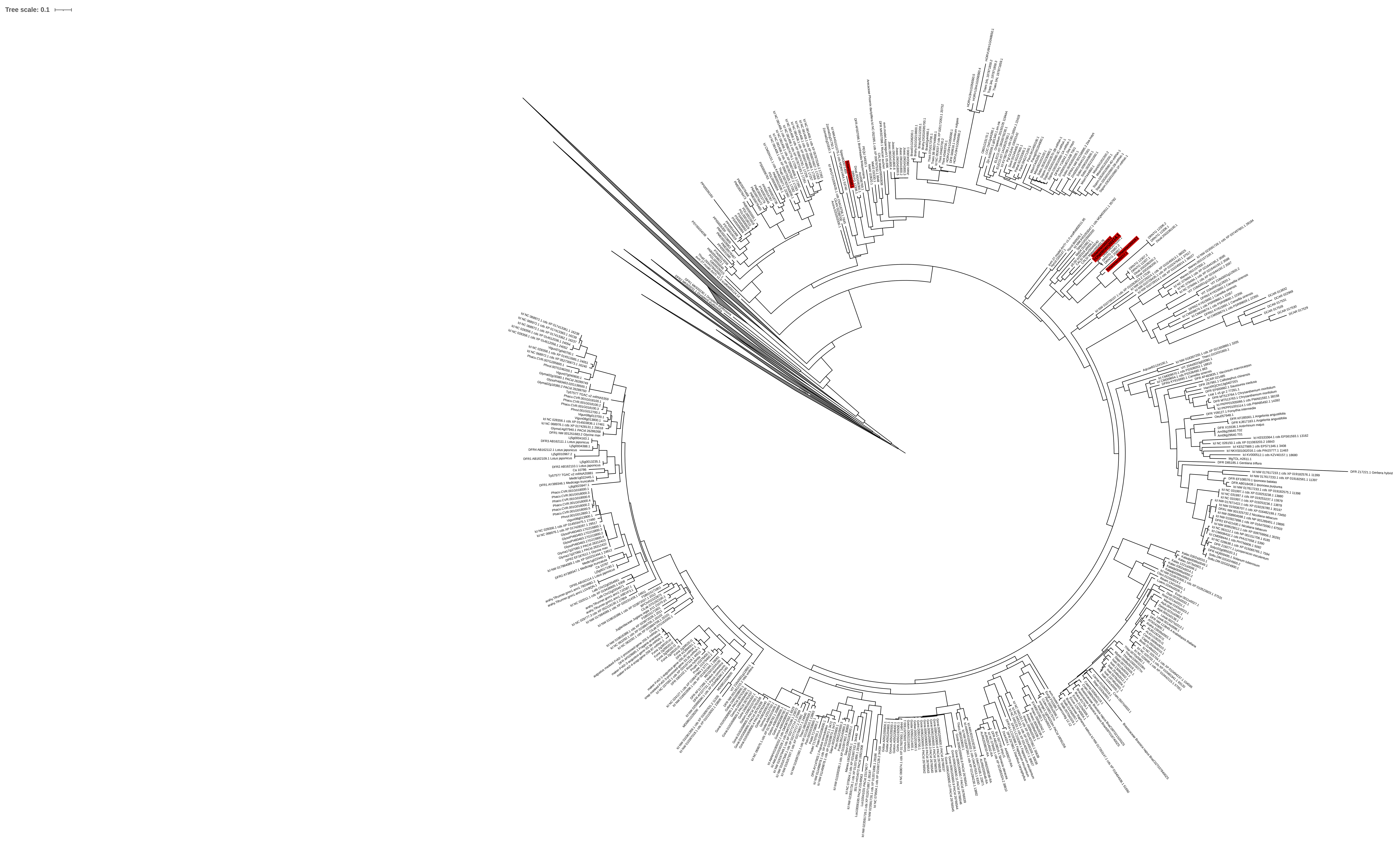
